# The heterochronic activation of TGF-β signaling drives the diversity of the avian sterna

**DOI:** 10.1101/2025.09.26.678725

**Authors:** Seung June Kwon, Zhaonan Zou, Mizuki Honda, Shiro Egawa, Shinya Oki, Yuji Atsuta

**Author notes:** Correspondence (Y.A.).

## Abstract

Adaptive evolution has driven the diversification of vertebrate skeletal morphology, enabling a wide array of locomotor modes and lifestyles. In volant birds (carinates), the sternum develops a ventral keel that provides an attachment site for massive pectoral muscles essential for powered flight. In contrast, flightless ratites have lost both flight capability and the keel, resulting in a striking morphological contrast between their sterna. Here, we interrogated the cellular and molecular basis underlying this morphological divergence by using chicken embryos as a carinate model and emu embryos as a ratite model. Through a series of analyses including spatiotemporal transcriptomics and a spheroid culture system, we found that TGF-β signaling, which promotes proliferation of ventral sternal chondroprogenitors, is activated in both species until the stage when the left and right sternal progenitors meet; however, in chicken this activation persists beyond this stage, driving ventral extension of the keel primordium, whereas in emu it shuts off early, culminating in the absence of a protruding keel. Together, our findings suggest that skeletal morphological changes associated with behavioral transitions can arise from heterochrony in developmental signaling, thereby deepening our understanding of the evolutionary logic shaping skeletal diversification.

## Introduction

The rich diversity of vertebrate forms and functions is fundamentally shaped by variation in bone morphology, which underlies differences in movement, feeding, and body structure. The skeletal template of vertebrates is established during embryogenesis through intramembranous ossification in the head and endochondral ossification in the trunk. Thus, with the exception certain postnatal skeletal modifications such as those associated with secondary sexual characteristics, interspecific diversity in skeletal morphology arises primarily during embryonic development^1,2,3^. Nevertheless, due in part to the limited availability of suitable experimental models, the mechanisms governing skeletal morphological diversification remain incompletely understood.

Among vertebrates, birds have undergone an extensive adaptive radiation. They have successfully expanded their ecological niches, including habitat range and dietary preferences, primarily through diversification of their skeletal morphology^4^. Carinates, the largest morphological group of modern birds, possess a skeletal system optimized for powered flight^5^. One of the most prominent features of their skeleton is the blade-like projection on the midline of the sternum, known as the keel. This keel serves as the attachment site for the massive pectoral muscles that generate the tremendous power required for flight. Consequently, keel height correlates with flight ability^6^; for example, hummingbirds, which exhibit exceptionally high flight capacity, have a relatively large keel^7^. Moreover, anatomical and fossil records suggest a correlation between acquisition and enlargement of the avian sternum and the evolution of flight ability^6,8^. The fact that other powered fliers, such as bats and extinct pterosaurs, also possess keel-like projections on their sterna further underscores the importance of the keel for flight^9,10^. In contrast, ratites—the other major group of modern birds—have lost both flight and the keel^11,12^. These cursorial, running-adapted birds including emus and ostriches, have flat sterna with only thin layers of pectoral muscles attached, which is consistent with their flightlessness.

The cellular and molecular mechanisms that determine the sternal morphological difference between carinates and ratites, namely the presence or absence of the keel, remain unknown. Previously, studies investigating the genetic mechanisms underlying sternum formation have primarily utilized conditional knockout mice. In the somatic lateral plate mesoderm (LPM), from which sternal progenitors (SPs) arise, the transcription factor *Prrx1* is expressed. By combining a *Prrx1* promoter that drives expression in the LPM with Cre recombinase^13,14^, SP-specific gene knockout (KO) can be achieved. Studies utilizing these conditional KO mice have identified the transcription factors *Runx1, Runx2*, *Sox9*, and *Tbx5* as being required for sternum formation^7,15,16^. However, Runx1/2 and Sox9 are master regulators of osteogenesis and chondrogenesis, respectively, and are essential for the formation of nearly all bones and cartilage. Moreover, because *Tbx5* is expressed in the sternum-forming regions of both carinate (chicken) and ratite (emu)^7,17^, these genes are unlikely to account for the evolutionary divergence in keel development.

In this study, we aimed to elucidate the differences in cellular behaviors during sternum cartilage formation between chicken and emu embryos, as well as to identify the molecular mechanisms that regulate these behaviors. The chicken is a traditional model organism for developmental studies, with established techniques for embryonic surgical experiments, such as tissue or bead transplantation, and for stable gene manipulation via *in ovo* electroporation using the transposon system^18^. Furthermore, a wealth of genome and gene expression resources is available for chicken^19^. On the other hand, the emu, which diverged from its most recent common ancestor with chickens around 110 million years ago (presumed to have been volant with a keeled sternum)^20^, has become an increasingly valuable model in the field of evolutionary developmental biology. This is partly because established embryonic staging is available^21^, and a chromosome-level genome assembly has been completed^22^, enabling detailed comparative studies^17,23^. First, we traced the specification timing, behavior and maturation state of SPs in chicken and emu embryos to investigate how differences in their dynamics contribute to keel formation. By performing gene expression profiling including spatial transcriptomics and *in vitro/in vivo* functional analyses, we find that prolonged activation of TGF-β signaling in ventral tip SPs maintains these cells in an immature, proliferative state in chicken but not in emu, a heterochrony that underlies the keel outgrowth. Our study provides evidence that heterochronic activation of a developmental signaling pathway, which arose during adaptive evolution, has driven the diversification of avian sternal morphology that underlies distinct locomotor strategies.

## Results

### The process of sternum formation shows no major differences between chicken and emu up to stage 34

In adults, the keel, which serves as the attachment site for the pectoral muscles, is present in chicken but absent in emu (Fig. 1a, b; Supplementary Movies 1 and 2), and this difference is already apparent by embryonic stage (St.) 36^7,21^. We therefore investigated whether differences occur in the behavior of the SPs between the two species prior to St. 36, specifically regarding the timing of their specification within the somatopleural layer and subsequent ventral migration (Fig. 1c-f).

**FIGURE 1.**
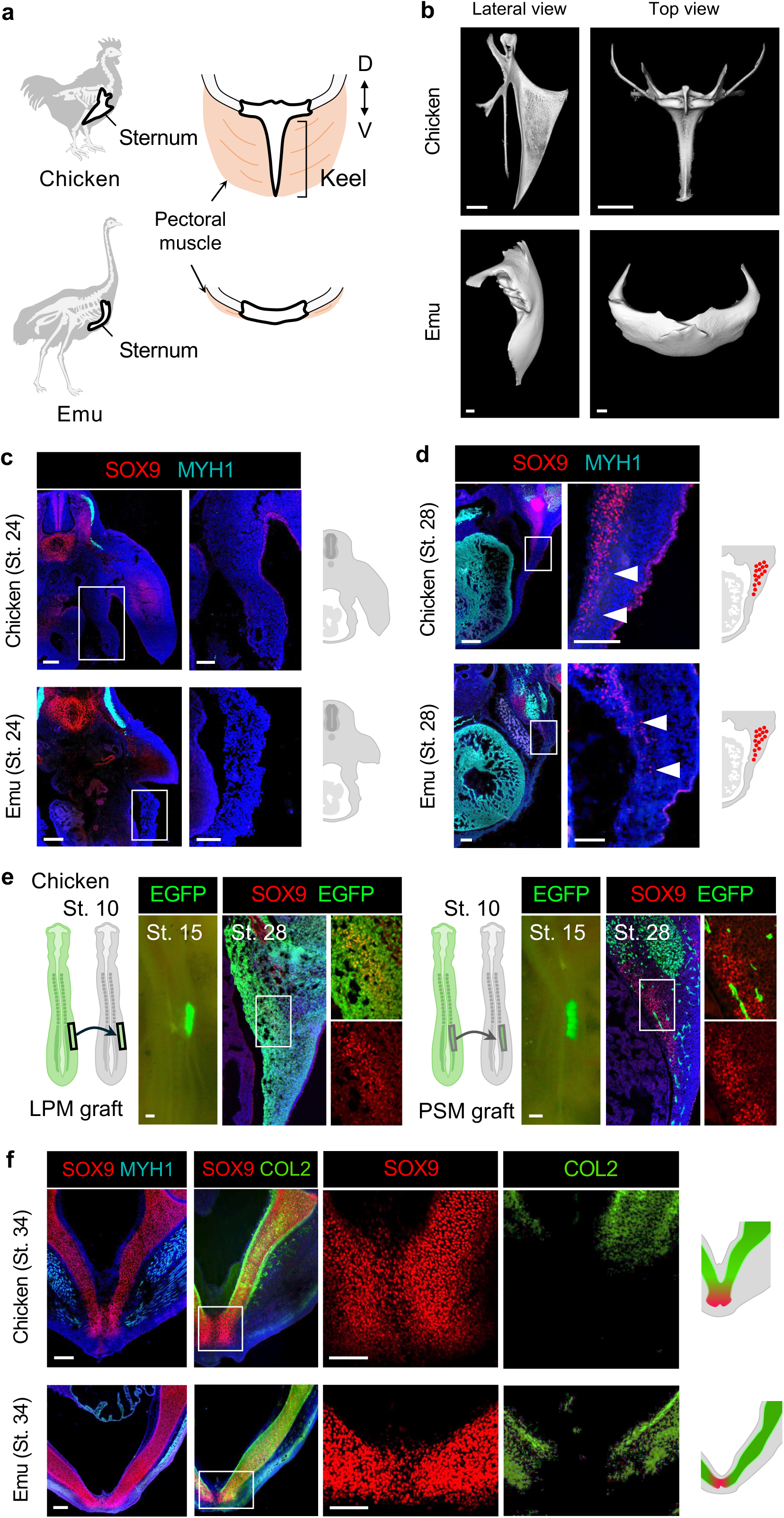
Comparable timing of sternum progenitor (SP) specification, ventral migration, and midline fusion in chicken and emu. **a** Schematics for thoracic regions of chicken and emu. The chicken sternum bears a midline projection called the keel, providing an attachment site for the large pectoral muscles (the pectoralis major and supracoracoideus muscles). The emu sternum lacks this projection, and its pectoral muscles are thin. **b** Lateral and top views of the three-dimensionally rendered sterna from an adult chicken and emu. **c**, **d** Transverse sections of the prospective sternal region in chicken and emu embryos at stage (St.) 24 (**c**) and 28 (**d**). Early chondrocytes are detected with an anti-SOX9 antibody (red), and muscle cells are detected with an anti-MYH1 antibody (cyan). At St. 28, SOX9-positive cells appear in both chicken and emu (arrowheads). **e** Reciprocal transplantation of chicken lateral plate mesoderm (LPM) or presomitic mesoderm (PSM). Tissues from St. 10 EGFP donor embryos were exchanged with the corresponding tissues of St. 10 wild-type host embryos. At St. 28, EGFP/SOX9 double-positive cells were observed in EGFP-LPM grafted embryos, whereas the two signals did not merge in EGFP-PSM grafted embryos (n = 3 each). **f** Sections of chicken and emu embryos at St. 34. SOX9 (red) and MYH1 (cyan), or SOX9 and collagen type II (COL2, green), are co-immunostained. Nuclei in transverse sections were counterstained with DAPI (blue). Scale bars, 1 cm (**b**), 200 μm; inset 100 μm (**c** - **f**).

At St. 24, SOX9 protein, an immature chondrogenic progenitor marker, is detected in the limb buds outgrowing adjacent to the prospective sternum-forming domain, indicating the initiation of chondrogenesis (Fig. 1c). In contrast, in the sternum-forming domain, SOX9 expression is not yet detected in either species (Fig. 1c).

SOX9 signals emerge in the sternum-forming regions of both chicken and emu at St. 28 (Fig. 1d). Because somitic cells contributing to rib progenitors are migrating into this region at this stage, where they meet the LPM to establish the lateral somitic frontier^24^, these SOX9-positive cells may originate from somites rather than from the LPM. We therefore traced their origin using EGFP-expressing chicken embryos^25^. At the prospective forelimb level along the anterior-posterior axis of wild type host embryos, we replaced either the LPM or the presomitic mesoderm (PSM) with EGFP-expressing donor tissue and examined whether the transplanted cells differentiated into SOX9-positive cells at St. 28 (Fig. 1e). As a result, in EGFP-labeled LPM grafted embryos, SOX9-positive cells expressed EGFP, whereas in EGFP-labeled PSM grafted embryos, SOX9-positive cells were EGFP-negative, indicating that SOX9-positive cells in the sternum-forming region are derived from the LPM (Fig. 1e). At the later St. 35, EGFP-LPM cells were incorporated into the sternal region, whereas EGFP-PSM cells gave rise to pectoral muscle cells (Supplementary Fig. 1), indicating that lineage tracing using EGFP-chickens was appropriately performed.

At the more advanced St. 31, these SOX9-positive SPs migrate further ventrally (Supplementary Fig. 2), and by St. 34, the left and right SPs in both chicken and emu meet at the midline and begin to fuse (Fig. 1f). In addition to SOX9, we performed immunostaining for Collagen II (COL2), a marker of differentiated cartilage cells, and found that by St. 34, all SPs except those at the ventral tip had differentiated into mature chondrocytes in both species (Fig 1f). Thus, until St. 34, ventral tip SPs remain immature in both chicken and emu.

Taken together, these results suggest that the timing of SP specification, ventral migration, and the process of chondrogenic maturation show no major differences between chicken and emu up to St. 34, when the bilateral SPs connect and commence the formation of a single sternum cartilage.

### Proliferative SPs persist at the tip of the keel-forming region in chicken

At St. 36, as previously observed in skeletal preparations^7^, the SPs that have reached the midline in chicken actively form ventrally elongating protrusions (Fig. 2a). By contrast, such ventral elongating structures are lacking in emu embryos (Fig. 2a). Of note, double immunostaining for SOX9 and COL2 revealed that, as seen at St. 34, ventral SPs in chicken were SOX9^+^/COL2^-^, while those in emu were SOX9^+^/COL2^+^, indicating that the emu ventral SPs had progressed toward more mature chondrocytes (Fig. 2a). In chicken, upon staining for Collagen IX (COL9), another marker of mature cartilage, together with SOX9, the ventral SPs remained SOX9 single-positive at both St. 34 and 36, indicating that these cells are indeed immature chondrocytes (Supplementary Fig. 3).

**FIGURE 2.**
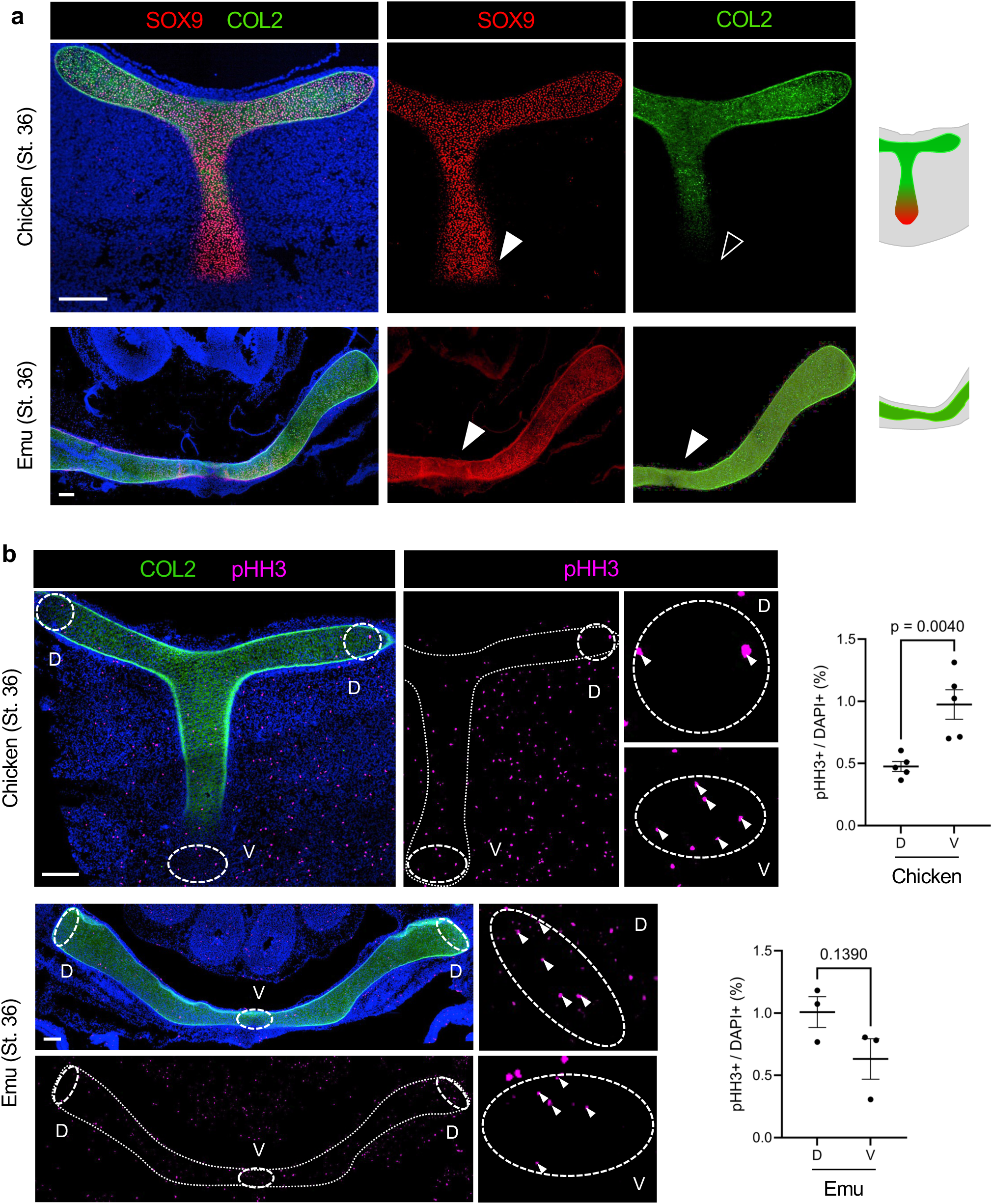
Ventral SPs mature earlier in emu than in chicken. **a** Transverse sections of St. 36 chicken and emu embryos. In chicken, sternal progenitors (SPs) located at the ventral tip of the keel-forming region are positive for SOX9 (red, arrowhead) but negative for COL2 (green, open arrowhead). All emu SPs are double positive for SOX9/COL2. **b** Mitotic cells in the sternal cartilage of St. 36 chicken and emu were detected using an antibody against phosphor-histone H3 (pHH3, magenta), and the mitotic cell ratio (pHH3^+^/DAPI^+^) was compared between the dorsal and ventral regions (arrowheads). Chicken samples: n = 15 views from 5 embryos (a total of 13,194 cells analyzed); emu samples: n = 9 views from 3 embryos (a total of 10,735 cells analyzed). The numbers shown in the plots indicate *p*-values, which were obtained using an unpaired two-tailed t-test. Scale bars, 200 μm.

Chondrocytes generally reduce their proliferative activity as they undergo maturation and hypertrophy^26^. Thus, in chicken, highly proliferative ventral SPs appear to contribute to the extension of the prospective keel region, whereas in emu, the reduced proliferative activity of ventral SPs may underlie the lack of keel extension. To test this possibility, we compared the proportion of dividing cells between ventral and dorsal SPs in chicken and emu, using phosphorylated histone H3 (pHH3) as a marker (Fig. 2b). We found that the proportion of pHH3-positive cells was higher in ventral SPs than in dorsal SPs in chicken (Fig. 2b). On the other hand, in emu, no difference in proliferation rate was observed between the dorsoventral regions, likely due to the lower proliferative capacity of differentiated ventral SPs (Fig. 2b).

These findings suggest that ventral SPs in chicken are maintained in an immature state until later developmental stages, and that keel extension is driven by these proliferative cells. Furthermore, these results imply that signaling pathways responsible for maintaining SPs in an immature state are persistently and specifically activated in chicken ventral SPs throughout later embryonic stages.

### Spatial transcriptomics reveals the activation of TGF-β signaling in keel-forming ventral SPs

Next, to characterize the gene expression profiles of SPs, we carried out transcriptome analysis. To compare ventral and dorsal SPs, we turned to photo-isolation chemistry RNA-Seq (PIC-RNA-Seq)^27^, which enables extraction of mRNA information exclusively from UV-irradiated regions of frozen tissue sections (Fig. 3a), thereby allowing direct comparison of their transcriptional profiles. Although this was the first attempt to apply PIC-RNA-Seq to avian cells, the sequencing metrics including read counts, mapping rates, and UMI numbers demonstrated that the PIC-RNA-Seq procedure was successfully implemented (Supplementary Fig. 4). Consequently, we reliably detected an average of 9,417 genes in chicken samples and 10,199 genes in emu samples (Supplementary Fig. 4).

**FIGURE 3.**
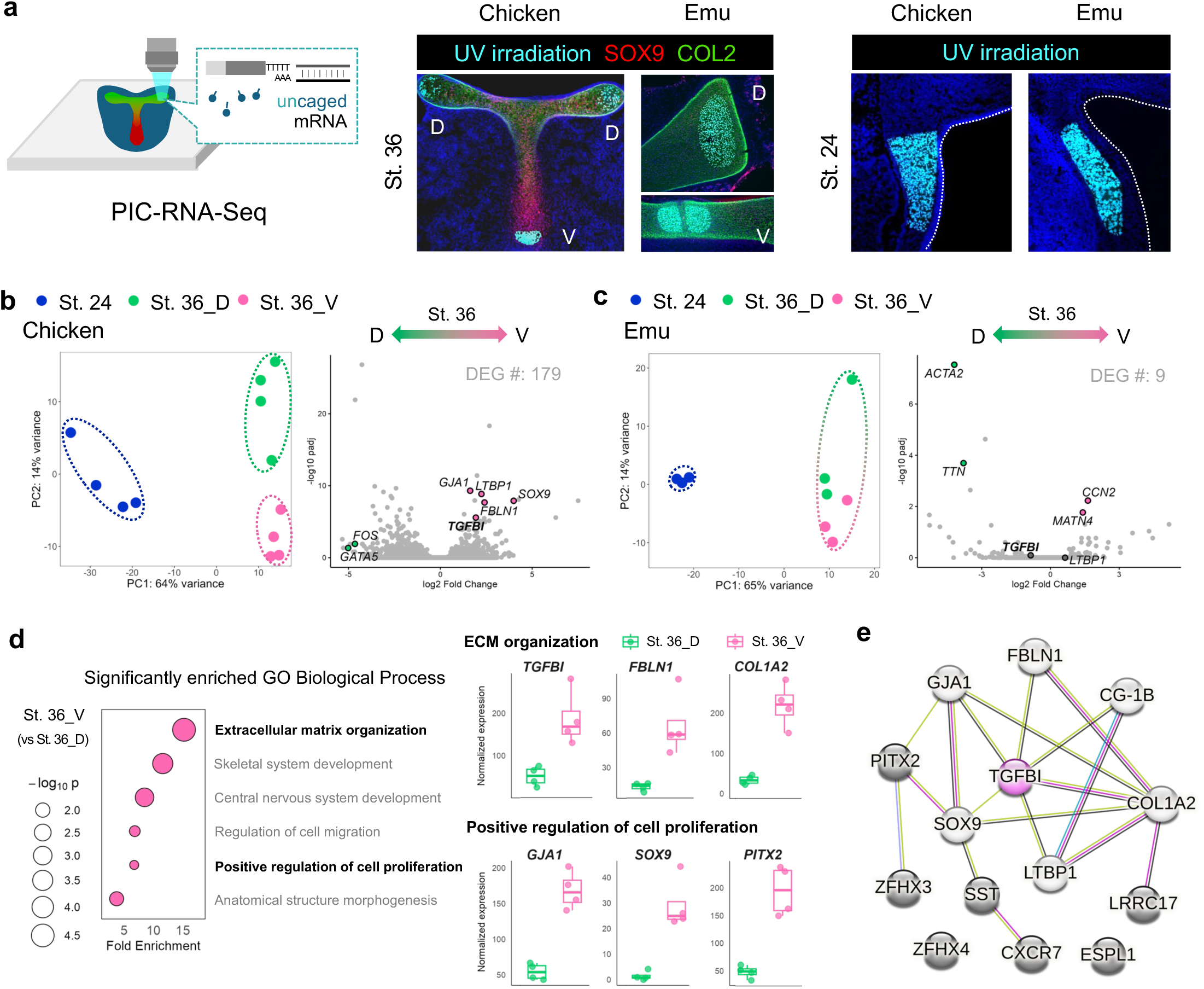
Specific activation of TGF-β signaling in ventral SPs during keel outgrowth. **a** For Photo-isolation chemistry (PIC)-RNA-Seq, the dorsal and ventral regions of the sternum were UV-irradiated (cyan) in St. 36 chicken and emu. As pre-chondrogenic samples, the SP-forming regions of St. 24 chicken and emu were also UV-irradiated. **b** PCA analysis showing clear separation between St. 24 and 36 chicken samples along the PC1 axis, while St. 36 dorsal and ventral samples separate along the PC2 axis. Volcano plot comparing St. 36 dorsal and ventral samples. Genes with values under threshold are colored gray. Among 179 differentially expressed genes (DEGs), representative ones are labeled. See also Supplementary Data 1 for additional DEGs. **c** PCA revealed that St. 24 and 36 emu samples are separated, whereas St. 36 dorsal and ventral samples show almost no separation. DESeq analysis detected only 9 DEGs between St. 36 dorsal and ventral samples. **d** Top 6 of Gene Ontology (GO) Biological Process terms enriched at St. 36 chicken ventral SPs, compared to St. 36 dorsal cells. See also Supplementary Data 4 for additional enriched GO terms. Box plots showing the differential expression of *TGFBI*, *FBLN1*, *COL1A2* (genes related to ECM organization), and *GJA1*, *SOX9*, *PITX2* (genes involved in cell proliferation). **e** Networks of genes upregulated in St. 36 chicken ventral SPs were described by STRING. Colors of lines (interactions) indicate text mining (green), experimentally determined (pink), co-expression (black), protein homology (violet), and curated database (blue). Colors of circles indicate TGFBI (pink), proteins with (white) or without (gray) direct interaction with TGFBI. Normalized expression and adjusted *p*-values are derived from DESeq2.

We conducted Principal Component Analysis (PCA) to compare the global patterns in gene expression among St. 24 and 36 SPs (Fig. 3b, c). Because cellular differentiation of SPs progresses from St. 24 to 36 in both chicken and emu, the samples at these two stages showed a clear separation (Fig. 3b, c). Focusing only on St. 36, a slight separation between dorsal and ventral SPs was observed along the PC2 axis in chicken (Fig. 3b), whereas no such dorsoventral difference was observed in emu (Fig. 3c). By analyzing differentially expressed genes (DEGs) between dorsal and ventral SPs at St. 36, we identified 179 DEGs in chicken and only 9 in emu (Fig. 3b, c; Supplementary Data 1). Among the chicken DEGs, in addition to *SOX9*, *TGF-β-Induced* (*TGFBI*) and *Latent TGF-β-Binding Protein 1* (*LTBP1*) were enriched in the ventral SPs (Fig. 3b). The transcription of *TGFBI* encoding an extracellular matrix protein is upregulated by TGF-β signaling^28,29^, and LTBP1 is known to regulate the secretion and activation of TGF-β proteins^30,31^. Thus, these data suggest that TGF-β signaling becomes activated in the ventral SPs during keel outgrowth. Notably, in emu, significant dorsoventral variations in the expression levels of these genes were not detected (Fig. 3c).

To decipher and compare the significant biological processes in chicken SPs, we performed Gene Ontology (GO) analysis, taking advantage of PANTHER and the chicken GO annotation file^32,33^. Several GO biological processes were enriched in the ventral SPs, including ones related to extracellular matrix organization and cell proliferation (Fig. 3d). Among the genes constituting these two biological processes, *TGFBI*, *Fibulin1*(*FBLN1*), *Collagen1A2* (*COL1A2*), *GJA1* (also known as *connexin43*), *SOX9*, and *PITX2* were particularly high at the ventral SPs (Fig. 3d). RNA-Seq data sets of chicken SPs were further subjected to STRING analysis^34^, to visualize inferred protein-protein association networks that were altered in the ventral SPs. Intriguingly, this revealed a network centered on TGFBI and LTBP1, which also included FBLN1, GJA1, COL1A2, and PITX2 (Fig. 3e). These data suggest that the activation of TGF-β signaling is closely associated with extracellular matrix remodeling (through TGFBI, LTBP1, FBLN1, and COL1A2), intracellular communication (via GJA1), and developmental regulation (via PITX2), implying that TGF-β positively regulates the organization and proliferation of ventral SPs through these pathways.

To summarize, our transcriptomic analysis raises the possibility that at St. 36, proliferation in chicken ventral SPs is maintained through the activation of TGF-β signaling, while in emu ventral SPs, the lack of TGF-β signaling activation is linked to reduced proliferative capacity.

### The activation of TGF-β signaling in ventral SPs terminates at an earlier developmental stage in emu than in chicken

Subsequently, we examined the dynamics of TGF-β signaling activation in SPs using the expression of *TGFBI* as a proxy. In addition to *TGFBI*, we assessed the expression of *LTBP1* and *TGF-β2*, which encode a mediator of TGF-β signaling and a TGF-β ligand, respectively, by *in situ* hybridization (Fig. 4).

**FIGURE 4.**
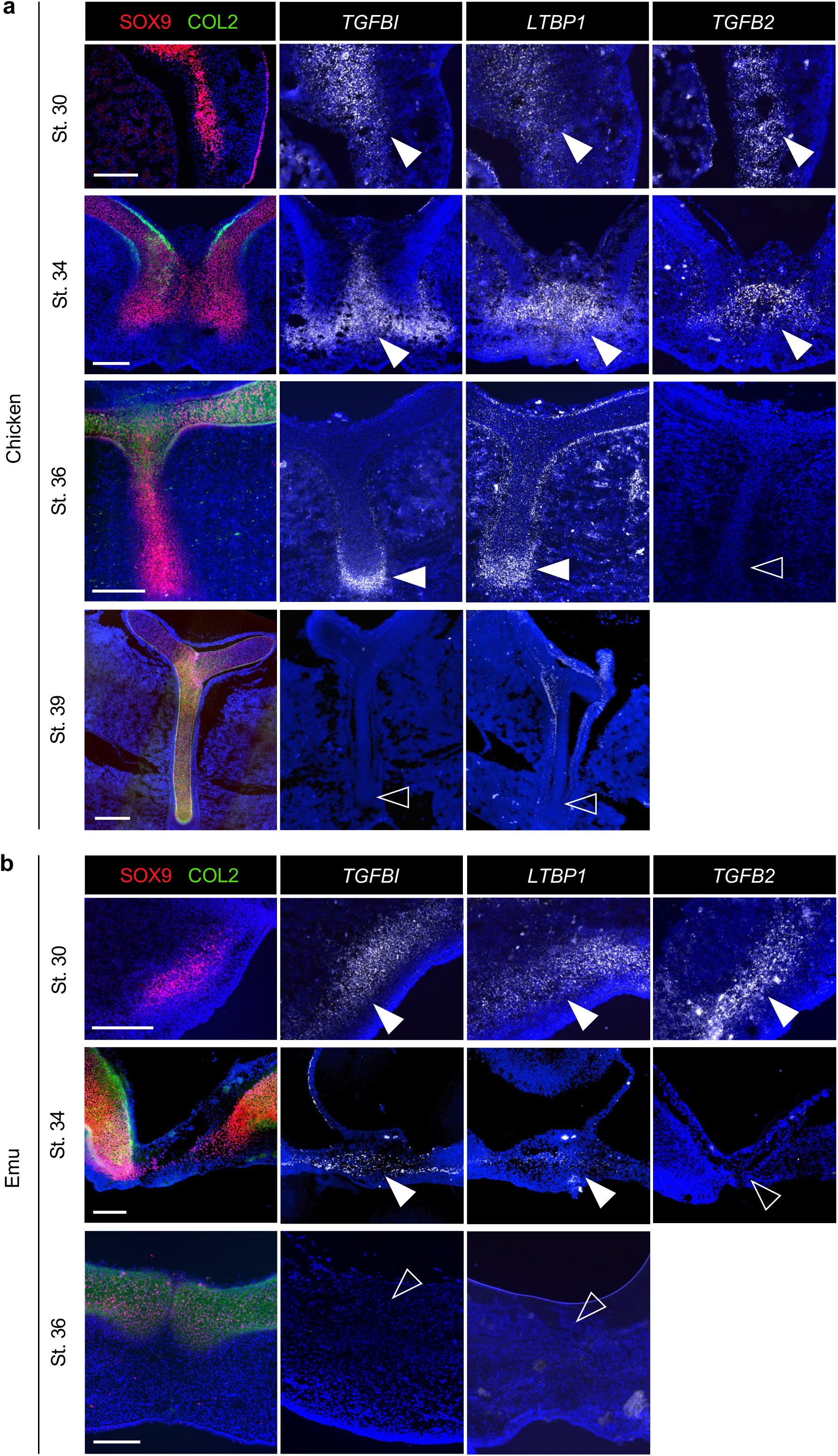
Heterochrony in TGF-β signaling activation between chicken and emu ventral SPs. **a** Fluorescent *in situ* hybridization (FISH) was used to detect the mRNA expression of *TGFBI*, *LTBP1*, and *TGFβ2* in transverse sections of chicken embryos from St. 30 to 39. At St. 30 and 34, all three genes were expressed in the ventral SP regions (arrowheads). At St. 36, *TGFBI* and *LTBP1* were detected in SOX9 (red) single-positive SPs or in their vicinity (arrowheads), whereas *TGFβ2* was not detected (open arrowhead). At St. 39, none of the three genes were detected. **b** FISH for emu samples. At St.30, all three genes were expressed (arrowheads), whereas at St. 34 *TGFβ2* is not detected (open arrowhead), and the expression of all three genes is absent at St. 36. Scale bars, 200 μm.

In chicken embryos, at St. 30 when SOX9 proteins are clearly observed in the sternum-forming regions, all three mRNAs were expressed in the regions (Fig. 4a). At St. 34, the expression of these genes was localized to the ventral SPs and the surrounding cells (Fig. 4a; Supplementary Fig. 5a). Co-staining of SOX9 and *TGF-β2* showed overlapping signals in the ventral SPs, implying that TGF-β signaling in these cells in an autocrine manner (Supplementary Fig. 5b). Consistent with the PIC-RNA-Seq results, the expression of *TGFBI* and *LTBP1* was detected in the ventral SPs at St. 36, whereas *TGF-β2* expression was no longer present at this stage (Fig. 4a; Supplementary Fig. 5a). At St. 39, when keel extension is nearly complete, none of these mRNAs were detected (Fig. 4a).

For *in situ* hybridization with emu embryos, we cloned partial cDNAs of emu *TGFBI*, *LTBP1*, and *TGF-β2*, and generated antisense riboprobes using these sequences as templates (Supplementary Data 2). These probes detected each respective mRNA in the expected locations, *i.e.*, *TGFBI* and *LTBP1* in the somites and *TGF-β2* in the limb mesenchyme^35,36^ (Supplementary Fig. 5c), thereby confirming that the probes we synthesized functioned properly. In emu embryos as well, mRNA expression of these three genes was detected in the sternal area at St. 30 (Fig. 4b). At St. 34, *TGFBI* and *LTBP1* were still detected in the ventral SPs. However, similar to the results in chicken at St. 36, *TGF-β2* expression had already disappeared by St. 34 (Fig. 4b). At St. 36, *TGFBI* and *LTBP1* were undetectable (Fig. 4b), indicating that TGF-β signaling was inactive in emu SPs at this developmental point.

Together with the PIC-RNA-Seq data, the *in situ* hybridization results indicate that TGF-β signaling is activated in the sternal-forming regions of both species until St. 34. However, whereas this activation is maintained over a longer developmental window in chicken, it terminates earlier in emu, and such heterochronic differences in activation may underlie the presence or absence of the keel.

### TGF-β signaling promotes the growth of both chicken and emu SP spheroids

To test whether TGF-β signaling indeed regulates the proliferation of chicken and emu SPs, we conducted *in vitro* assays. To this end, we applied a three-dimensional culture system, originally developed for the induction and maintenance of chondrocytes from human pluripotent stem cells^37^, to chicken and emu SPs (Fig. 5a). Under this chondrogenic medium and three-dimensional culture conditions, LPM cells isolated from St. 24 chicken or emu embryos aggregated and formed spheroids composed of SOX9-positive cells (Fig. 5a, b). Although this culture system does not fully reproduce *in vivo* sternal development, we cultured the spheroids for 15 days, as emu embryos require approximately 14 days to progress from St. 24 to 36^21^.

**FIGURE 5.**
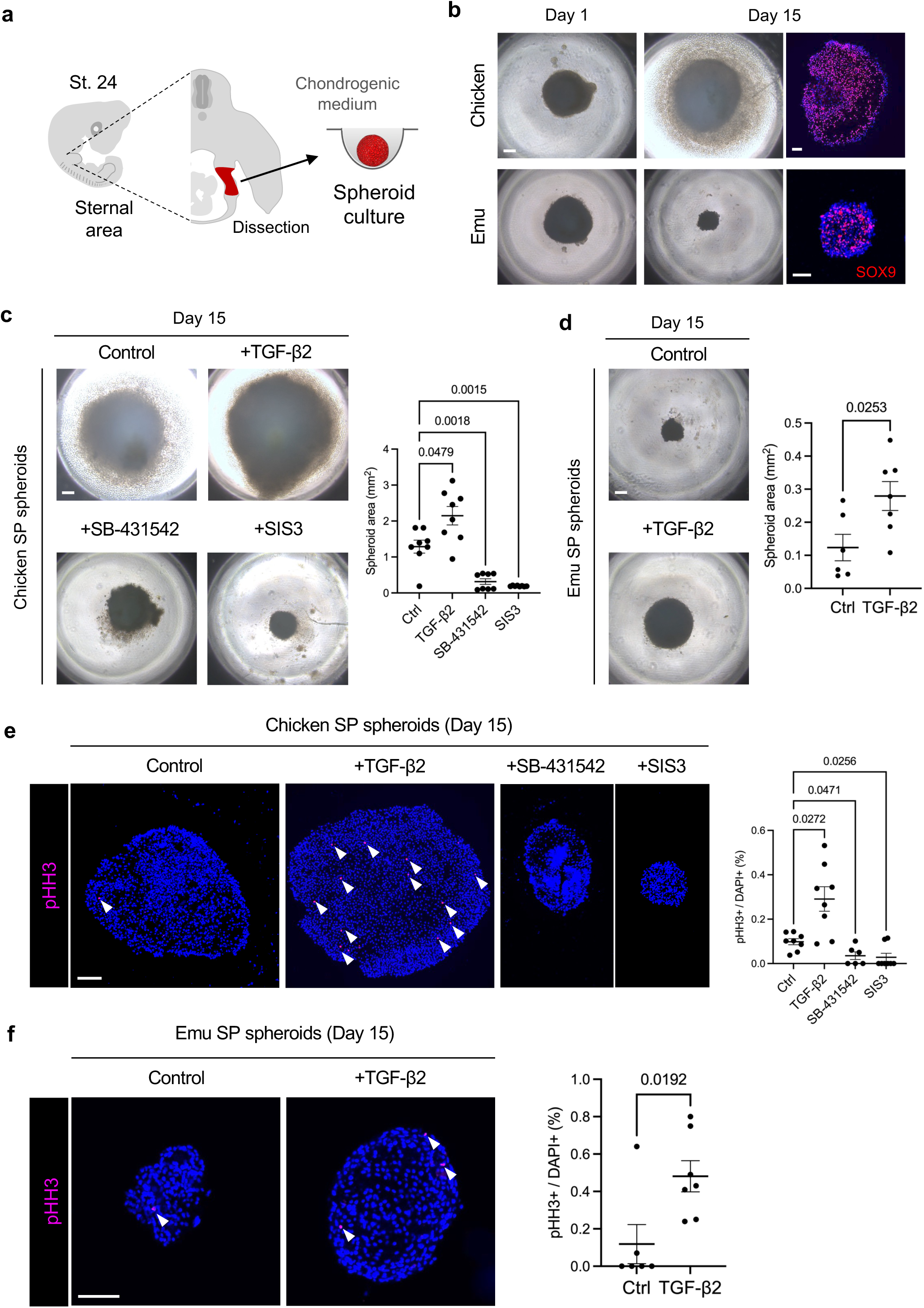
TGF-β signaling is necessary and sufficient to stimulate SP proliferation. **a** Schematic of spheroid culture of SPs. **b** Three-dimensionally cultured SPs derived from St. 24 chicken and emu. After 15 days of culture, the SPs become SOX9-positive (red). **c** Chicken SP spheroids cultured with DMSO (control), TGF-β2 protein, SB-431542 (a TGF-β inhibitor), or SIS3 (a specific Smad3 blocker). Spheroid areas in the images were measured (n = 8 each). **d** Emu SP spheroids cultured with DMSO (control) or TGF-β2 protein (n = 6 and 7). **e** Chicken SP spheroids cultured for 15 days were stained with an antibody against pHH3, and the mitotic cell ratio (pHH3^+^/DAPI^+^) was measured (n = 8, 8, 6 and 8). **f** Likewise, the mitotic cell ratio (pHH3^+^/DAPI^+^) of emu SP spheroids cultured for 15 days was measured (n = 6 and 7). *p*-values in the plots of (**c**) and (**e**) were obtained using Brown-Forsythe and Welch ANOVA and Dunnett’s T3 multiple comparison. *p*-values in (**d**) and (**f**) were obtained using an unpaired two-tailed t-test. Scale bars, 200 μm (**b**; bright-field, **c**, **d**), 75 μm (**b**; SOX9, **f**), 150 μm (**e**).

When TGF-β2 protein was added to the basal medium to augment intracellular TGF-β signaling, both chicken and emu spheroids became significantly larger than the control group, as expected (Fig. 5c, d; Supplementary Fig. 6). In contrast, treatment of chicken SP spheroids with the TGF-β inhibitors, SB-431542 or SIS3^38,39^, resulted in a marked reduction in spheroid size (Fig. 5c; Supplementary Fig. 6). To further investigate whether TGF-β activation promotes cell division, we measured the proportion of mitotic cells within the spheroids using pHH3 as a marker. In chicken SP spheroids treated with TGF-β2, the proportion of pHH3-positive cells was approximately threefold higher compared with the control group (Fig. 5e). Conversely, treatment with TGF-β inhibitors led to a dramatic reduction in the number of pHH3-positive cells (Fig. 5e). In emu SPs, administration of TGF-β ligand similarly resulted in a significant increase in the proportion of pHH3-positive cells, as observed in chicken SPs (Fig. 5f). These results strongly suggest that TGF-β signaling is both necessary and sufficient for the proliferation of avian SPs.

### TGF-β signaling is essential for keel formation in chicken embryos

We then interrogate the possible role of TGF-β signaling in sternal formation *in vivo*. Owing to the difficulty of obtaining fertilized emu eggs in sufficient numbers and the absence of a keel in the emu, we examined the *in vivo* role of TGF-β in sternal formation exclusively in chicken embryos. To ask whether TGF-β signaling is required for the outgrowth of sternal-forming regions, we set out to graft SB-431542-soaked beads into the coelom adjacent to the prospective sternal region of St. 15 chicken embryos (Fig. 6a). This manipulation cannot continuously inhibit TGF-β signaling in SPs; nevertheless, the absence of *TGFBI* expression in the sternal-forming region at St. 24, two days after bead implantation, indicates that the onset of TGF-β signaling activation was at least delayed (Fig. 6b). As a result of the bead implantation experiments, ventral elongation of the sternal-forming region was not disturbed in St. 32 embryos grafted with DMSO-soaked beads (Fig. 6c). On the other hand, in embryos grafted with SB-431542-soaked beads, impairment of ventral extension in the sternal-forming region was found on the implanted side (Fig. 6c). At the later St. 36, as at St. 32, keel elongation was diminished on the SB-431542-soaked beads-grafted side, whereas no effect was observed with DMSO-soaked control beads (Fig. 6d). Together, these findings indicate that TGF-β signaling is essential for proper sternum and keel development.

**FIGURE 6.**
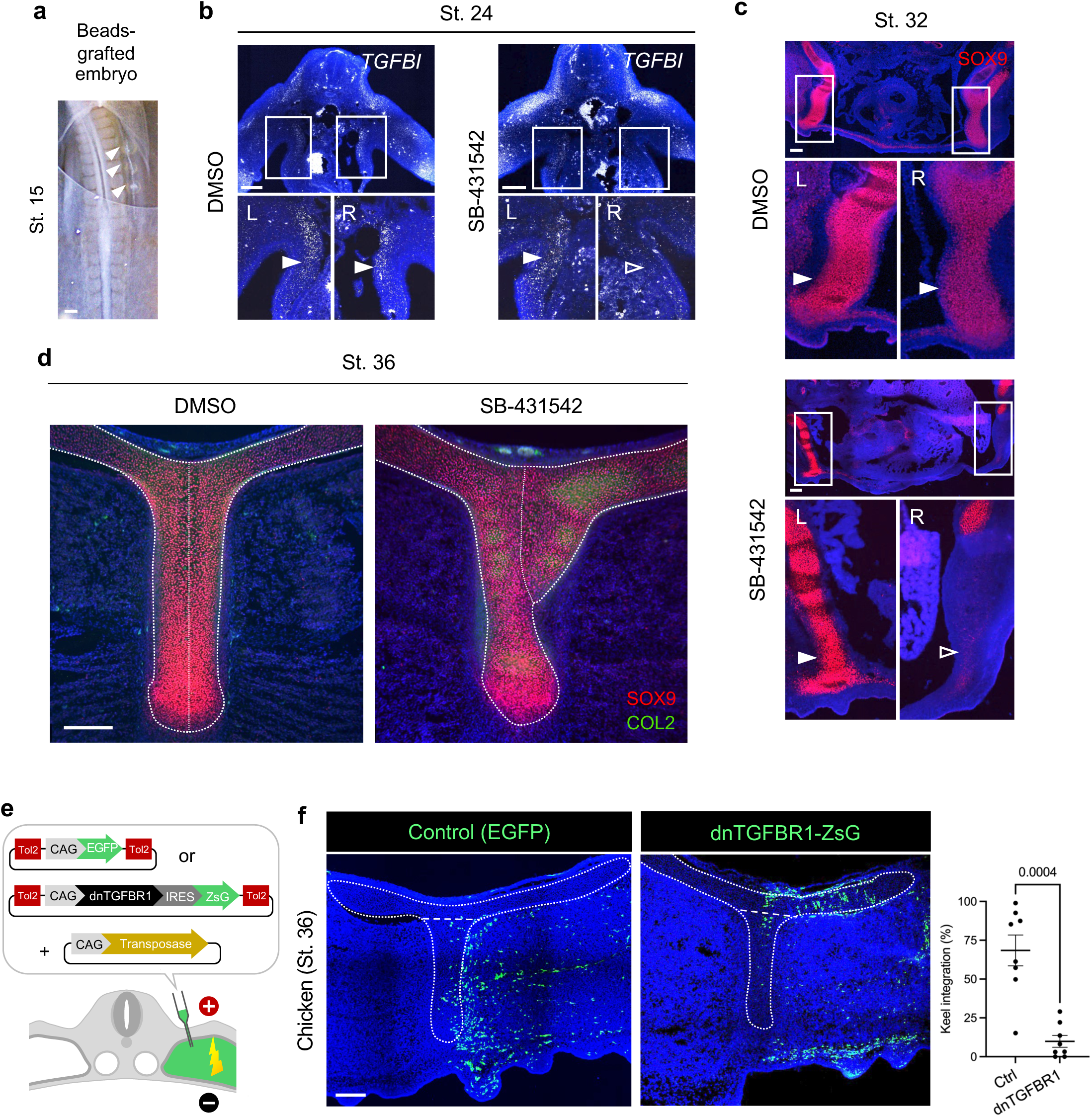
Requirement of TGF-β signaling for keel extension. **a** Three beads soaked in DMSO (control) or SB-431542 were grafted into the right coelom of St. 15 chicken embryos (arrowheads). **b** FISH of St. 24 embryos with bead grafts. *TGFBI* expression was diminished in the LPM on the bead-grafted side (open arrowhead). **c** At St. 32, no abnormalities were observed in the ventral extension of SPs in control embryos (n = 0/7), whereas ventral extension was inhibited in SB-bead grafted embryos (open arrowhead; n = 4/9). **d** At St. 36, control embryos formed a symmetrical keel as seen in normal embryos, and no abnormal phenotype was observed (n = 0/9), whereas in SB-bead grafted embryos, the keel on the bead-grafted side was shortened, resulting in an asymmetric keel (n = 3/10). **e** Plasmids used for electroporation into the somatopleural layer. dnTGFBR1 and ZsG denote a dominant-negative form of TGF-β receptor I and ZsGreen1, respectively. The negative electrode was positioned ventrally and the positive one dorsally, without direct contact with the embryo. **f** In St. 36 chicken embryos, dnTGFBR1-expressing cells are less frequently distributed in the keel region of the sternal cartilage compared with control EGFP-expressing cells. *p*-value in the plot of (**f**) was obtained using an unpaired two-tailed t-test. Scale bars, 200 μm.

Next, we attempted to genetically inhibit TGF-β signaling by *in ovo* electroporation, introducing a dominant-negative form of TGF-β receptor 1 (dnTGFBR1) into SPs. This dnTGFBR1 harbors a lysine-to-arginine substitution, abolishing kinase activity toward Smad2/3^40^. To achieve sustained expression of the dominant-negative receptor within SPs and hence suppress TGF-β signaling, we employed the Tol2 transposon system^18^ (Fig. 6e). As a control, expression of EGFP in SPs resulted in approximately 70% of EGFP-positive cells being integrated within the keel region on average (Fig. 6e). In contrast, dnTGFBR1-expressing cells failed to distribute to the keel region and instead were largely located in the dorsal part of the sternal cartilage (Fig. 6e). These findings suggest that SPs require cell-autonomous activation of TGF-β signaling in order to contribute to the keel-forming region.

These *in vivo* experiments demonstrate that TGF-β signaling regulates the behavior of ventral SPs, thereby playing a pivotal role in chicken keel formation.

## Discussion

Elucidating the mechanisms underlying skeletal morphological diversification is a central objective in Evo-Devo biology. In this study, we focused on the sternum, which displays a simple yet striking morphological difference among birds, and demonstrate that diversification of the avian sternal shape is, at least in part, attributable to a heterochronic shit of TGF-β signaling activation in the ventral tip of the sternum (Fig. 7). Before the keel elongation starts in the chicken, overall sternal development is comparable between the chicken and emu: (1) the chondrogenic specification and ventral fusion occur around the same developmental stage; (2) early chondrocytes (SOX9-positive and COL2-neagtive) are maintained in the ventral edge of the sterna; and (3) activation of TGF-β signaling is observed in ventral SPs of both species up to the ventral fusion stage, and this activation promotes the proliferation of SPs. However, in emu, ventral SPs mature earlier than in chicken and form flat sternum, while in chickens they persist to remain and through active proliferation, contribute to the keel extension.

**FIGURE 7.**
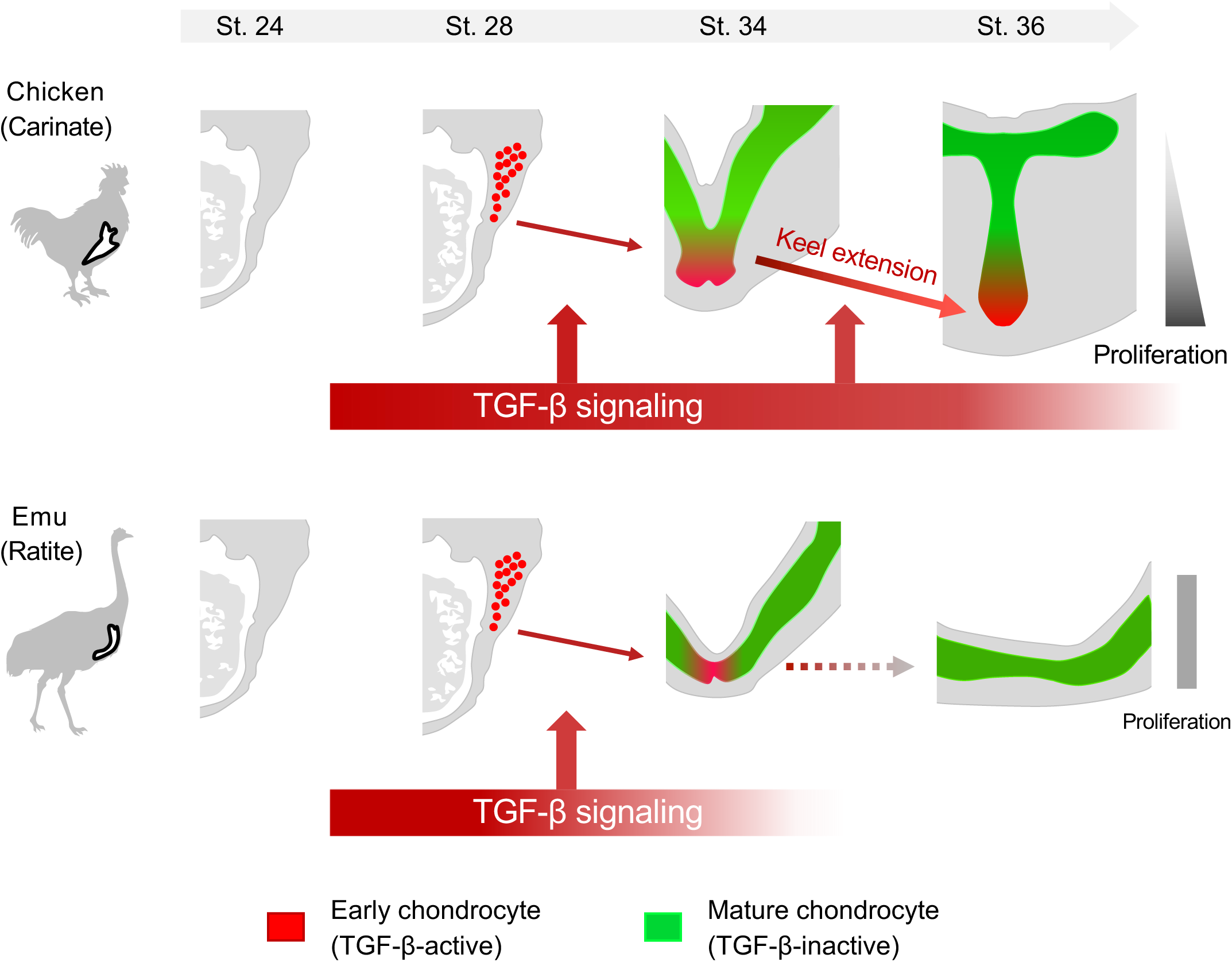
Schematic model illustrating the cellular and molecular mechanisms underlying avian sternal diversity. In chickens, SOX9-positive SPs arise within the somatopleural layer of the LPM at St. 28. Ventral SPs actively proliferate by receiving TGF-β2, secreted either by themselves or by neighboring cells, thereby activating TGF-β signaling and driving ventral extension of the sternal cartilage. This TGF-β signaling activity continues until St. 36, during which the keel-forming region elongates. In emus, SPs are specified at St. 28 in a manner similar to chickens, and until St. 34 the sternal cartilage extends toward the ventral midline under the influence of TGF-β signaling. However, unlike in chickens, TGF-β signaling activation has terminated by St. 36, and consequently keel formation does not occur.

### SP specification in the somatic LPM

While it was previously known that the SPs of chicken originate from the somatic LPM at the forelimb level^7^, the precise area where and when the specified SPs emerge had not been documented. Using immunostaining for SOX9, we found that chondrogenic specification of SPs occurs in the ventral region of somatic LPM at approximately St. 28 in both chicken and emu (Fig. 1). Given that LPM-specific *Sox9* deletion results in complete loss of the sternum in mice, the SOX9 expression in the somatic LPM, which specifies SPs, represents an initial and crucial step in sternum formation in vertebrates. Interestingly, unlike in birds and mice, snake embryos, which lack a sternum, show no Sox9 expression in the somatic LPM (Supplementary Fig. 7), suggesting that sternum formation fails due to the absence of SP specification. Thus, the difference in sternum presence versus absence, rather than in its morphological diversity, may have been determined during evolution by whether the mechanism specifying SPs was maintained or lost.

The molecular entity controlling SP specification remains unknown. According to our PIC-RNA-Seq data (Supplementary Data 3), expression of *TGF-*β*2* is detected in the sternum-forming region as early as St. 24, prior to St. 28 when SOX9 protein expression becomes evident. Moreover, the fact that LPM-specific *TGFBR2*-KO and *TGF-*β*2 null* mice exhibit a bifurcated sternum, phenocopying *Sox9* mutants, further supports TGF-β signaling as a strong candidate^16,41,42^. In addition, SPs cultured in chondrogenic media containing BMP4 differentiated into SOX9-positive cells even in the absence of exogenous TGF-β supplementation (Fig. 6b), suggesting -similar to chondrogenic specification in limb mesenchyme^43^ – that BMP signaling may also play a central role in SP specification.

### Role of TGF-β signaling in sternal development

TGF-β signaling mediates diverse biological processes, including cell proliferation, migration, differentiation, and extracellular matrix organization during embryogenesis^44,45,46^. In the context of chondrogenesis, TGF-β signaling is known to promote proliferation of chondrocytes and inhibit terminal differentiation, among other regulatory roles^47,48^. Conditional *TGFBR2*-KO mice display shortened limbs together with sternal defects, primarily due to reduced chondrocyte proliferation^41^. These mice also exhibit an expanded hypertrophic zone in the growth plate, in part because of accelerated chondrocyte maturation, and cultured limb mesenchyme from these mice shows enhanced differentiation^41^. Collectively, these findings suggest that TGF-β signaling promotes proliferation while restraining maturation of chondrocytes derived from the somatic LPM. Intriguingly, conditional knockout of *TGFBR2* in *Col2a1*-expressing cells does not affect the sternum in mice^49^, consistent with the absence of TGF-β activity in COL2-positive SPs observed in our study (Fig. 4). Extending previous findings, we further provide evidence for a role of TGF-β signaling in regulating the sternal chondrogenesis by enhancing the growth of ventral immature SPs.

In addition to its role in proliferation, TGF-β signaling may also contribute to SP migration. Following specification, the left and right SPs migrate from the dorsal to ventral direction, eventually fusing at the ventral midline. During this migration, ventral-specific activation of TGF-β signaling was observed in immature SPs and their neighboring cells. These ventral leading cells express the ligand *TGF-*β*2*, which has been shown to promote the migration of a mesenchymal precursor cell line^50^. In addition, enhanced expression of *TGFBI* has been reported to promote migration in glioma cells^51^. Ventral leading cells in chicken express *TGFBI* throughout the ventral fusion and keel elongation, implying that their ventral localization may determine the direction of keel extension.

The involvement of TGF-β signaling in sternal development appears to be conserved in humans, as mutations in related genes have been linked to congenital chest wall deformities. Pectus excavatum is the most common chest deformity (about 1 in 300-1,000 births^52,53^) where the sternum grows inward and affects the function of heart and lung. The second most is pectus carinatum, where the sternum protrudes outward and causes chest pain and disturbs respiration. Despite their relatively high prevalence, the causative genes for these congenital disorders have not been clearly identified yet. However, because some patients with Marfan or Loeys-Dietz syndrome, in whom pectus excavatum is one of the clinical manifestations, carry mutations in either *TGFBR1* or *TGFBR2*^53,54,55^, it is possible that misregulation of TGF-β signaling contributes to the development of pectus excavatum. In addition, patients homozygous for a missense mutation affecting a highly conserved amino acid of *Fibulin-4* (*FBLN4*) exhibited pectus excavatum^56,57^. Fibulin-4 deficiency is known to elevate TGF-β2 levels and thereby enhance TGF-β signaling^57,58^. Although the pathogenesis of congenital chest deformities remains unclear, the overgrowth of sternocostal chondrocytes has been proposed as one of the major contributing factors^59^. Given that our findings demonstrate a role for TGF-β in promoting SP proliferation, it is plausible that excessive TGF-β signaling contributes to hyperplastic chondrocyte growth in the thoracic region, ultimately leading to chest abnormalities. Further investigation on avian sterna may identify regulatory mechanisms that sustain TGF-β2 expression in chickens, which could provide valuable insight into the developmental basis of congenital chest wall deformities.

### Regulatory mechanisms underlying the heterochronic expression of *TGF-*β*2*

Our FISH results revealed the early termination of *TGF-*β*2* expression in emu ventral SPs. While this heterochronic expression may result from various several evolutionary scenarios, one possibility could be species-specific differences in transcriptional regulation. The *TGF-*β*2* promoter contains a cAMP-response element (CRE) site bound by the transcription factor CREB^60,61^, and an enhancer of *TGF-*β*2* is known to be bound by NF-κB^62^. Our PIC-RNA-Seq data indicate that CREBs, their coactivator CREBBP, and NF-κBs are expressed in both chicken and emu SPs (Supplementary Data 3). These findings suggests that future investigations into cis- and trans-regulatory divergence, incorporating genomes of species with proportionally large sterna such as hummingbirds as well as snakes that entirely lack a sternum, may provide deeper insight into the evolutionary diversification of sternal development.

### Heterochronic signaling as a driver of morphological diversification

Although it is evident that many morphological differences among species originate during embryonic development, surprisingly few studies have directly implicated heterochrony in developmental signaling as a proximate cause of interspecific variation. Among the rare examples, Shapiro et al. (2003) provided compelling evidence that heterochronic termination of *Sonic hedgehog* expression contributes to digit number variation in skink lizards, highlighting the potential temporal shifts in signaling to drive morphological change^63^. Yet, such studies have largely remained descriptive, without experimental validation of the causal role of signaling heterochrony. Our work extends this line of inquiry by demonstrating that the persistence of TGF-β signaling differs between chicken and emu, and by functionally showing that TGF-β plays an essential role in promoting the proliferation of chicken SPs required for keel extension. By linking a heterochronic shift in signaling activity to a specific cellular mechanism that underlies a clear morphological divergence, our findings move beyond descriptive correlations and advance our understanding of the developmental processes that generate skeletal diversity.

### Limitations of this study

The major limitations of this study are twofold. First, it remains technically challenging to generate chickens with KO mutations in either TGF-β receptors or the TGF-β signaling mediators Smad2/3, and therefore we were unable to demonstrate whether global genetic ablation of TGF-β signaling shortens the keel in chicken. Currently, KO chickens are mainly produced through transplantation of cultured KO-primordial germ cells (PGCs). However, because maintenance of PGCs requires TGF-β signaling^64^, it is not feasible to establish PGCs with KO mutations in TGF-β signaling genes. Although the generation of *Prrx1*-CreERT and TGFBR-floxed chicken lines for conditional KO would be theoretically possible, it would require an extraordinary amount of time. Second, due to the limited availability of emu embryos, we were unable to perform sufficient *in vivo* functional analyses, let alone generate transgenic animals with prolonged TGF-β signaling.

## Methods

### Animals

Fertilized white leghorn chicken (*Gallus* gallus) eggs purchased from Yamagishi and Takeuchi poultry farm were incubated at 38.5 °C and staged according to Hamburger and Hamilton stage (St.)^65^. Fertilized emu (*Dromaius novaehollandiae*) eggs from Kiyama farm (Saga, Japan) were incubated and turned every 30 min at 36 °C, and staged as previously described^21^. In chicken embryos, embryonic day 1.5 (E1.5), E2, E4, E5.5, E6, E7, E8, E9, E10, and E13 correspond to St. 10, 15, 24, 28, 30, 32, 34, 35, 36, and 39, respectively. In emus, E5, E9, E13, E14, E20 and E23 correspond to St. 15, 24, 28, 30, 34, and 36, respectively. Fertilized Japanese striped snake (*Elaphe quadrivirgata*) eggs were obtained from Japan Snake Institute (Gunma, Japan) and were incubated at 28 °C until 6 days post oviposition, when the facial and visceral arch structures of the embryo become similar to chicken St. 30. All experiments using avian embryos were performed under ethical approval of Kyushu University (No. A25-248-0; A25-247-0).

### Three-dimensional (3-D) rendering of adult chicken and emu sterna

The original data of adult chicken and emu skeletons were obtained from the following URLs. 3-D images were generated using Amira^66^ by volume/surface rendering in a pseudo-color, then 2-D images were captured in orthographic views.

- *Gallus gallus* (FMNH B 492179; Media ID 000095954):

https://www.morphosource.org/concern/parent/000S26880/media/000095954

- *Dromaius novaehollandiae* (MZB 92-0121):

https://sketchfab.com/3d-models/dromaius-novaehollandiae-92-0121-esternum-f81f7db70591406eae92c9a3810b7873

### Immunohistochemistry

Embryos were dissected in phosphate-buffered saline (PBS) and fixed in 4% paraformaldehyde (PFA)/PBS at 4 °C overnight. Embryos were then rinsed with PBS and incubated overnight in 30% sucrose/PBS at 4°C. After another overnight incubation with 30% sucrose/PBS in OCT compound (1:2; Sakura Finetek), embryos were cryo-embedded in OCT. Sectioning was performed on a NX70 cryostat (Thermo-Fisher Scientific) at -10 to -15°C with 14 μm-thick sections attached to Frontier micro slides (Matsunami). Cryosections were rinsed in PBS to dissolve the OCT, and when immunostaining for collagens, antigen retrieval was performed by heating the vessel containing slides with 1x citrate buffer (GenoStaff, 10x G-Activate, pH 6). Otherwise, tissue permeabilization was done instead, by applying 0.5% Triton X-100 (Sigma) in PBS for 10 min at room temperature (RT). Next, slides were washed with 1x TNT buffer (0.1M Tris-HCl pH 7.5, 15 mM NaCl, 0.1% Tween20) for 5 min, and blocking was done with 1% blocking reagent (Roche) in 1x TNT for 45 min at RT. Incubation with primary antibodies was performed for overnight at 4°C. Primary antibodies used in this study are: rabbit anti-Sox9 for early chondrocytes (1:1000; Merck Millipore, AB5535), mouse anti-Myh1E for the myosin heavy chain (1:1500; DSHB, MF20), rabbit anti-pHH3 for mitosis (1:1000; Merck Millipore, 06-570), mouse anti-Col2a1 (DSHB, II-II6B3) and Col9a1 (DSHB, D1-9) for chondrocytes (1:500). After washing for 5 min in TNT 3 times, slides were incubated for 2 hours at 4°C with secondary antibodies: Alexa Fluor goat anti-rabbit 555 and donkey anti-mouse 488 (1:500; Thermo-Fisher). Slides were then washed for 5 min in TNT 3 times and mounted using Antifade Mounting Medium with DAPI (Vectashield). Images were obtained using a Dragonfly confocal microscope (Andor) or DM6000B (Leica).

### Cell counting and area measurement

For chicken and emu St. 36 sections, dorsal and ventral area of the sternum was decided as an ellipse with a major axis about 220 μm, then total cell numbers (DAPI) and proliferating cell numbers (pHH3) were counted by eye. For SP spheroids and electroporated St. 36 chicken sections, DAPI and pHH3 of fluorescent signals were counted (total 13,194 and 10,735 cells were counted for St. 36 chicken and emu sections, and 141,053 and 13,736 cells were counted for chicken and emu SP spheroids). Spheroids were imaged by CKX53 cell culture microscope (Olympus), and the area was measured using Fiji^67^.

### Tissue and beads transplantation experiment

Transplantation experiments using EGFP-expressing transgenic chicken embryos were performed at Harvard Medical School with the kind support of Dr. Clifford J. Tabin and under the guidelines of the Harvard Medical School Institutional Animal Care and Use Committee. Fertilized EGFP eggs were obtained from Clemson University^25^. From St. 10 EGFP embryos, the presumptive PSM corresponding to somites 15-20 (forelimb level) or the LPM located at the level of presumptive somites 15-20 was excised using a sharpened tungsten needle and transplanted into the corresponding region of wild-type embryos (Charles River) at the same stage (see also Fig. 1e).

For beads transplantation into St. 15 chicken embryos, AG1X2 ion-exchange beads (formate form; BioRad) were incubated in 100mM SB-431542 (Selleck) or DMSO (Nacalai Tesque) for 2 h at RT before grafting three beads into the right coelom^68^ of sternal region at the level of somites 15-18.

### Spatial transcriptomic analysis

Sternal area of St. 24 and St. 36 chicken and emu embryos were dissected in PBS, and embedded in OCT for 5 min at RT. Subsequently, tissues were placed into an aluminum cryomold, then the mold was immersed in isopentane/dry ice for flash-freezing and kept at - 80 °C. Cryosections at a thickness of 10 μm were mounted on Frontier micro slides and air-dried. Samples were first tested for proper RNA integrity number, then standard PIC-RNA-Seq with TE treatment was conducted as described^27,69^. Region of interest (ROI) for irradiating UV light on St. 24 samples was decided as the somatic LPM of sternal region, about 600 μm ventral from the end of the forelimb buds. For St. 36, after additional immunostaining of SOX9 and COL2 using same antibody and methods with Triton X as immunohistochemistry, dorsal and ventral ROI were decided as ellipses with a major axis about 220 μm. After in vitro transcription and library preparation as in Honda *et al.*^27^, quality of cDNA library was assessed using Bioanalyzer 2100 or TapeStation (Agilent Technologies) before sequencing on the Illumina NovaSeq 6000 platform (Illumina).

UMIs and barcodes in the reads were extracted using UMI-tools (ver. 1.0.0), then the reads were aligned to the reference genome obtained from Ensembl (chicken: GRCg6a, GCA_000002315.5; emu: ZJU2.0, GCA_016128335.2) using HISAT2^70^ (ver. 2.1.0). Read counts were quantified with featureCounts^71^ (ver. 2.0.3), and DESeq2^72^ (1.38.3) was used to perform the differential gene expression analysis and principal component analysis (PCA), then generate volcano plots. ggplot2^73^ (https://ggplot2.tidyverse.org.) and normalized counts obtained from DESeq2 were used for making box plots. GO analysis was conducted using PANTHER^33,74^ (ver. 18.0), and the top GO terms from the PANTHER hierarchy were represented in the plot. Protein-protein interaction network was analyzed with STRING (ver. 12.0)^75^, based on DEGs with an adjusted *p*-value < 5 x 10^-6^.

### Probe synthesis and *in situ* hybridization (ISH)

Total RNA was extracted from St. 19 chicken and emu embryos using Nucleospin RNA Plus (Macherey-Nagel). cDNA was synthesized by PrimeScript II 1^st^ strand cDNA synthesis kit (Takara) and used as a template for subsequent PCR. PCR products were subcloned into pBlueScript II SK (+) vector using Ligation high ver.2 (Toyobo). To synthesize anti-sense riboprobes, T3 or T7 RNA polymerase (Promega) and DIG RNA Labeling Mix (Roche) were used. Sequences of probes and primers used to amplify gene fragments are listed in Supplementary Data 2. To confirm the RNA probes, whole mount ISHs were performed as described^76^, using NBT/BCIP (Roche) for color reaction. Section fluorescent ISH was conducted as described previously^76^ with TSA Vivid 520 (Tocris Bioscience) diluted in RNAscope LS multiplex TSA buffer (1:1000; ACD) for about 20 min reaction.

### Sternal progenitor (SP) spheroid culture

Basal chondrogenic medium for SP spheroid culture was made as previously reported^37^ with 10 mM β-glycerophosphate (Wako), 10ng/mL BMP-4 (Nakalai Tesque) and 10 nM triiodothyronine (T3; Selleck) added.

Left and right sternal area of St. 24 chicken and emu embryos were dissected in PBS. Ectoderm was removed after a 10 min incubation in 0.2% Trypsin/EDTA (Wako) at RT using fine forceps. Dissected sternal tissues were then transferred into 15% FBS/DMEM in a 1.5 mL tube. After brief pipetting, the tube was centrifuged at 1500 rpm for 5 min at RT and the supernatant is removed. Cells were resuspended in chondrogenic medium added with 10 μM Y-27632 (Focus Biomolecules), and cultured in PrimeSurface 96V plate (Sumitomo Bakelite), one sternal area per well. From day 2, medium was changed to chondrogenic medium supplemented with 20 ng/mL TGF-β2 (Cosmo Bio) or TGF-β signaling inhibitors (5 μM SB-431542 or SIS3 HCl; Selleck). The medium was changed every 2-3 days.

### *In ovo* electroporation and expression vectors

The coding sequence (CDS) of a dominant negative form of TGFBR1 (dnTGFBR1) was amplified from pCMV5B-TGFBR1-K232R (Addgene plasmid #11763)^40^ by PCR using KOD One (Toyobo), and subcloned into pT2A-CAGGS-IRES2-ZsGreen1^77^ by Gibson Assembly Cloning Kit (New England Biolabs). The plasmid was sequence-verified before the electroporation, and primer sequences are listed in Supplementary Data 2. pT2A-CAGGS-EGFP and pCAGGS-T2TP were shared by Yoshiko Takahashi (Kyoto University)^18,78^. All DNA solution was prepared at 4 μg/μl and mixed with Fast Green FCF (Wako) before use for visualization.

For *in ovo* electroporation, DNA was injected into the right coelomic cavity of St. 14 chicken embryos using microcapillary pipettes with aspirator tube assemblies (Merck). The ne Three (+) electric poring pulses of 50 V, 0.5 ms, were given, followed by 8 (+/-) transfer pulses of 5 V, 10 ms, with 10 ms interval between pulses (Super Electroporator NEPA21-type II, NEPA GENE).

### Statistical analysis

All statistical analyses were performed with Prism 10.5 (GraphPad), and statistical details can be found in the legends. Measurements are represented as the mean ± SEM. No statistical method was used to predetermine the sample size. The experiments were not randomized, and the investigators were not blinded to allocation during experiments and outcome assessment.

## Acknowledgements

We greatly acknowledge Dr. Clifford J. Tabin (Harvard Medical School) for careful reading of the manuscript, as well as Drs. Daisuke Saito, Yoshiki Hayashi, Yoshinao Hosaka (Kyushu University), and Yoshiko Takahashi (Kyoto University) for helpful discussion. We also thank the Center for Advanced Instrumental and Educational Support of the Faculty of Agriculture (Kyushu University) and Transcriptomics Kenkyukai (General incorporated association, Fukuoka) for the use of Cryostar NX70 (Thermo Fisher Scientific) and RNA-Seq support, respectively. This work was supported by Research Support Project for Life Science and Drug Discovery (Basis for Supporting Innovative Drug Discovery and Life Science Research [BINDs]) from AMED under Grant Number JP23ama121017 and JP25ama121017. This study was also funded by the JST FOREST Program (Grant Number JPMJFR214G to Y.A.), JSPS KAKENHI (Grant Numbers JP25K09649 to Y.A. and JP24KJ1793 to S.J.K), The Sumitomo Foundation (to Y.A.), and The Takeda Science Foundation (to Y.A.).

## Author contributions

Conceptualization: Y.A.; Study design: S.J.K. and Y.A.; Methodology: S.J.K., Z.Z., M.H., S.E. and Y.A.; Investigation: S.J.K., Z.Z., S.O. and Y.A.; Visualization: S.J.K., Z.Z. and S.E.; Funding acquisition: S.J.K. and Y.A.; Writing-original draft: S.J.K. and Y.A.; All authors reviewed and approved the manuscript.

## Competing interests

The authors declare no competing interests.

## Figure legends

**Supplementary Figure 1.**
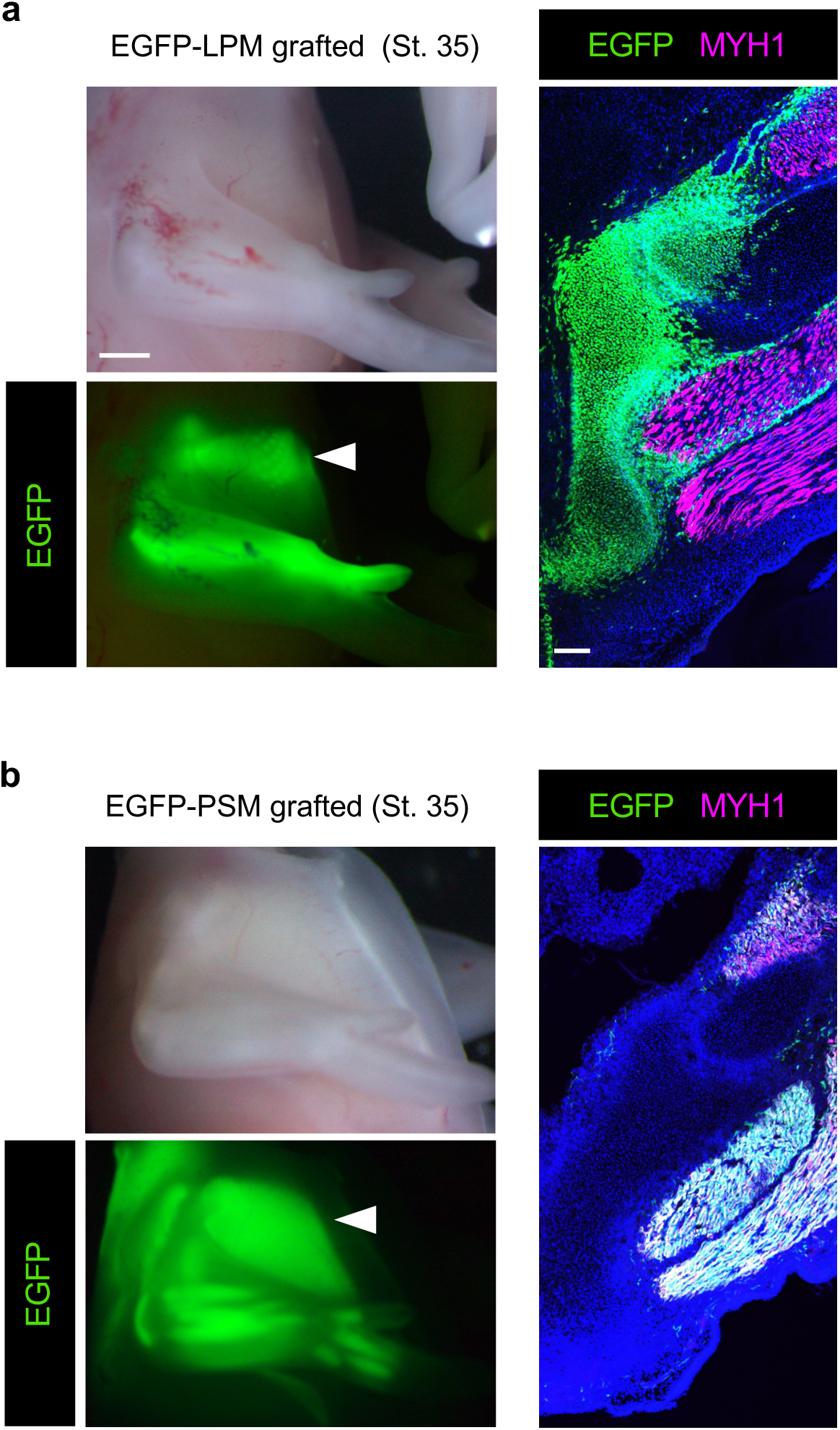
Grafted EGFP-lateral plate mesoderm (LPM) contributed to the sternal region, whereas grafted EGFP-presomitic mesoderm (PSM) gave rise to the pectoral muscles. **a** A whole-mount view and transverse section of a St. 35 embryo grafted with EGFP-labeled LPM (see also Fig. 1e). Keel-forming EGFP-positive cells are indicated by an arrowhead. EGFP signals do not merge with MYH1 (magenta). **b** A whole-mount view and transverse section of a St. 35 embryo grafted with EGFP-labeled PSM. The grafted EGFP-positive cells differentiated into MYH1-positive pectoral muscle cells. Scale bars, 1 mm, 100 μm.

**Supplementary Figure 2.**
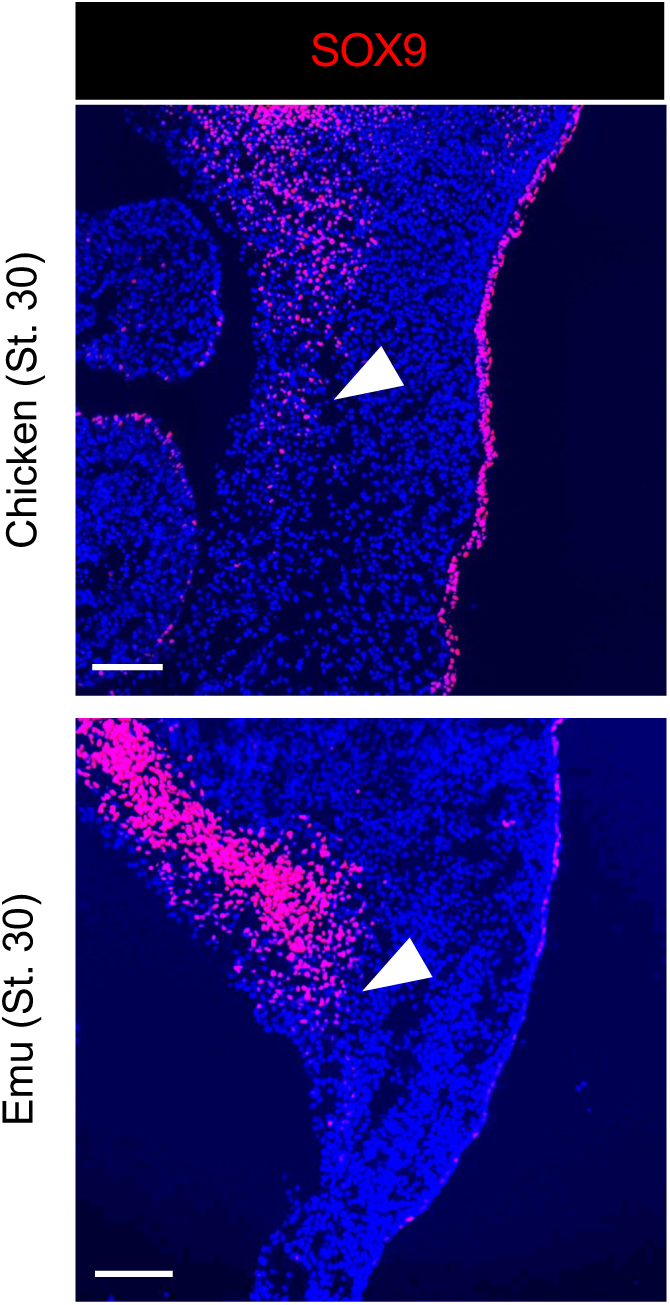
SOX9 signals in the sternum-forming region in St. 32 chicken and emu. Signals for SOX9 proteins (red) were detected at the sternum forming region of St. 32 chicken and emu embryos (arrowheads). Scale bars, 200 μm.

**Supplementary Figure 3.**
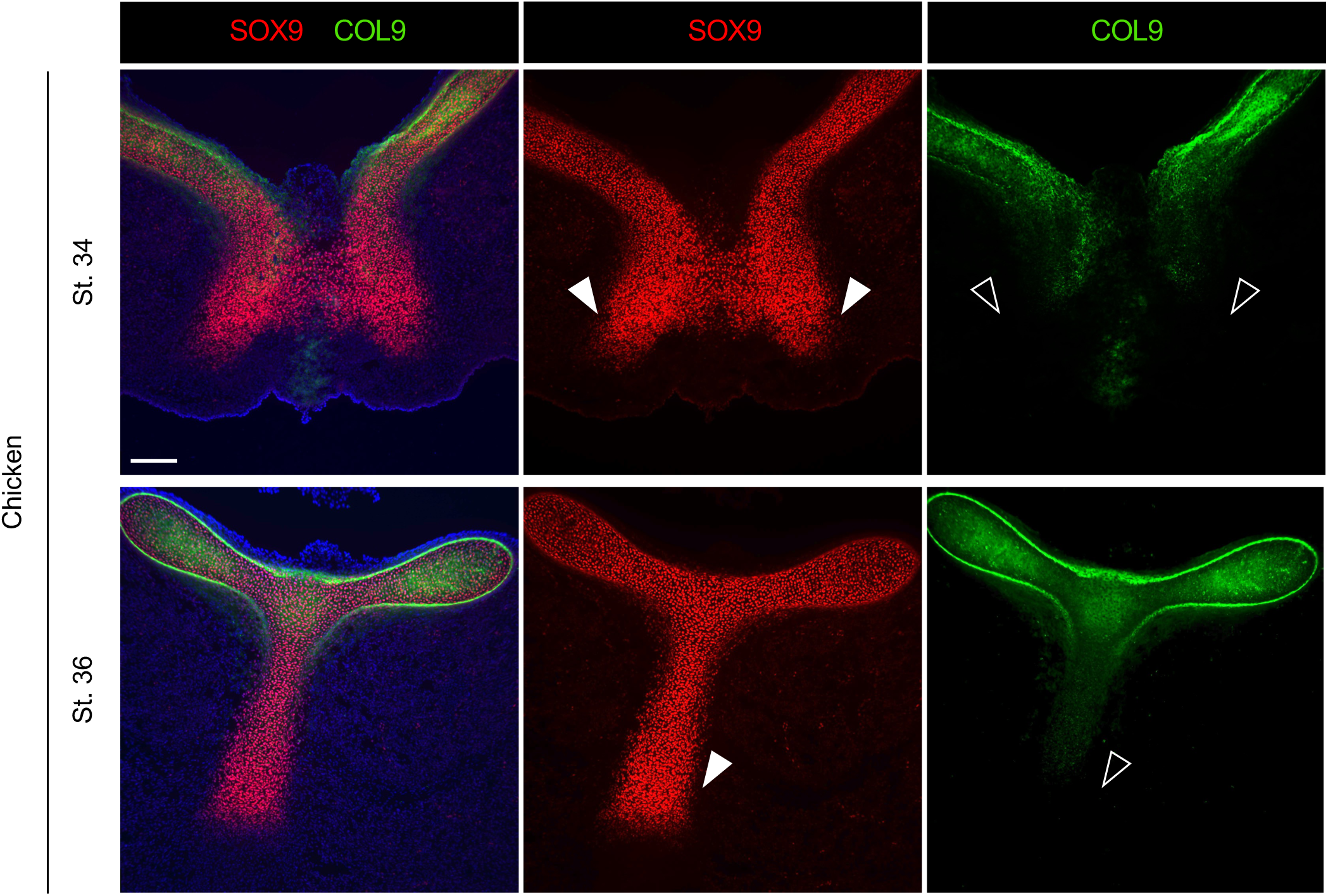
Dorsal SPs have differentiated into mature, COL9^+^ chondrocytes in St. 34 and 36 chickens. Sternum cartilage of St. 34 and 36 chickens immunostained with antibodies against SOX9 (red) and collagen IX (COL9, green). At both stages, ventral SPs were SOX9-positive and COL9-negative (arrowheads and open arrowheads). Scale bar, 150 μm.

**Supplementary Figure 4.**
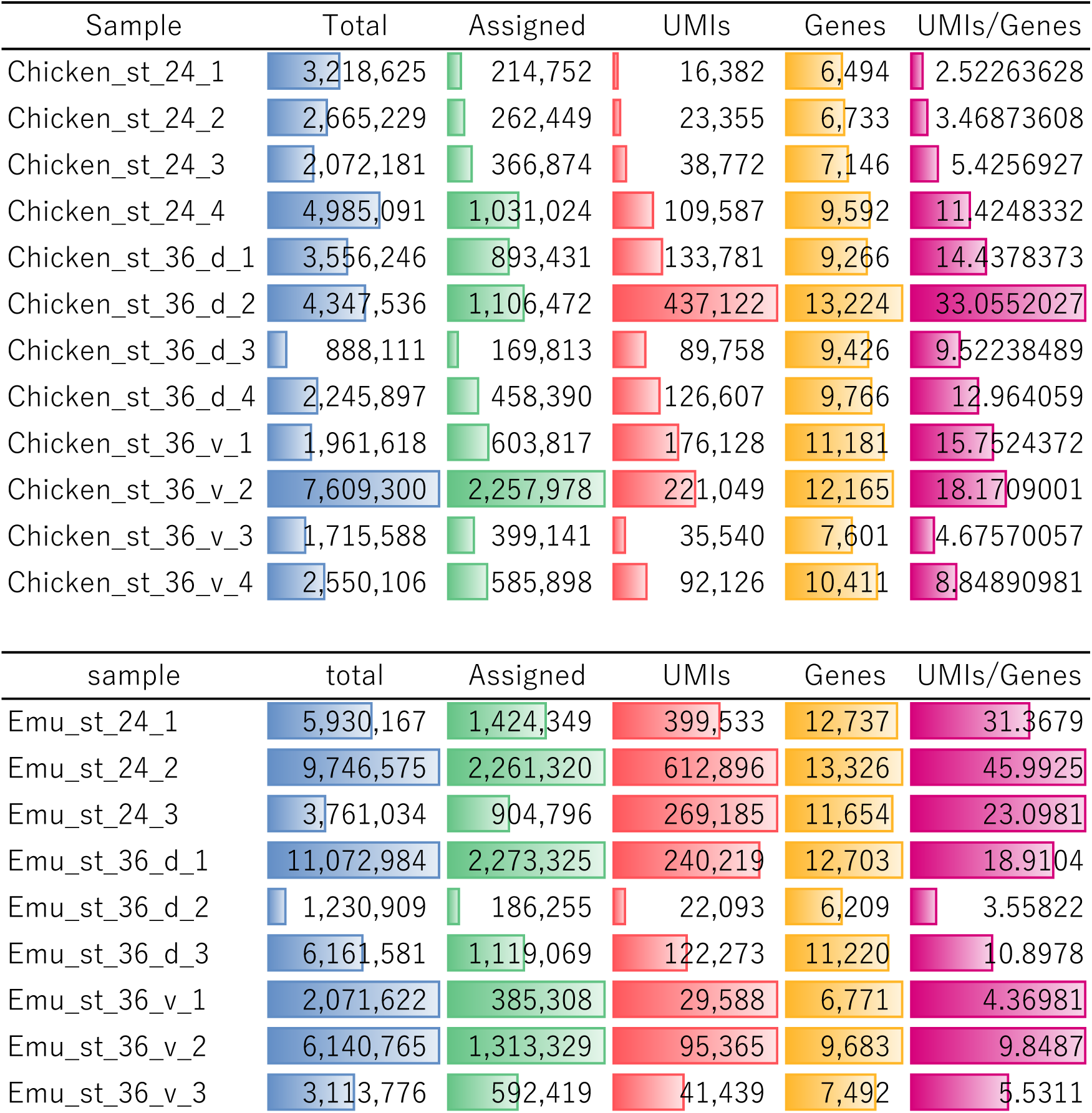
The sequencing metrics of PIC-RNA-Seq. Numbers of total reads, assigned reads, unique molecular identifiers (UMIs), and detected genes are shown for each PIC-RNA-Seq samples.

**Supplementary Figure 5.**
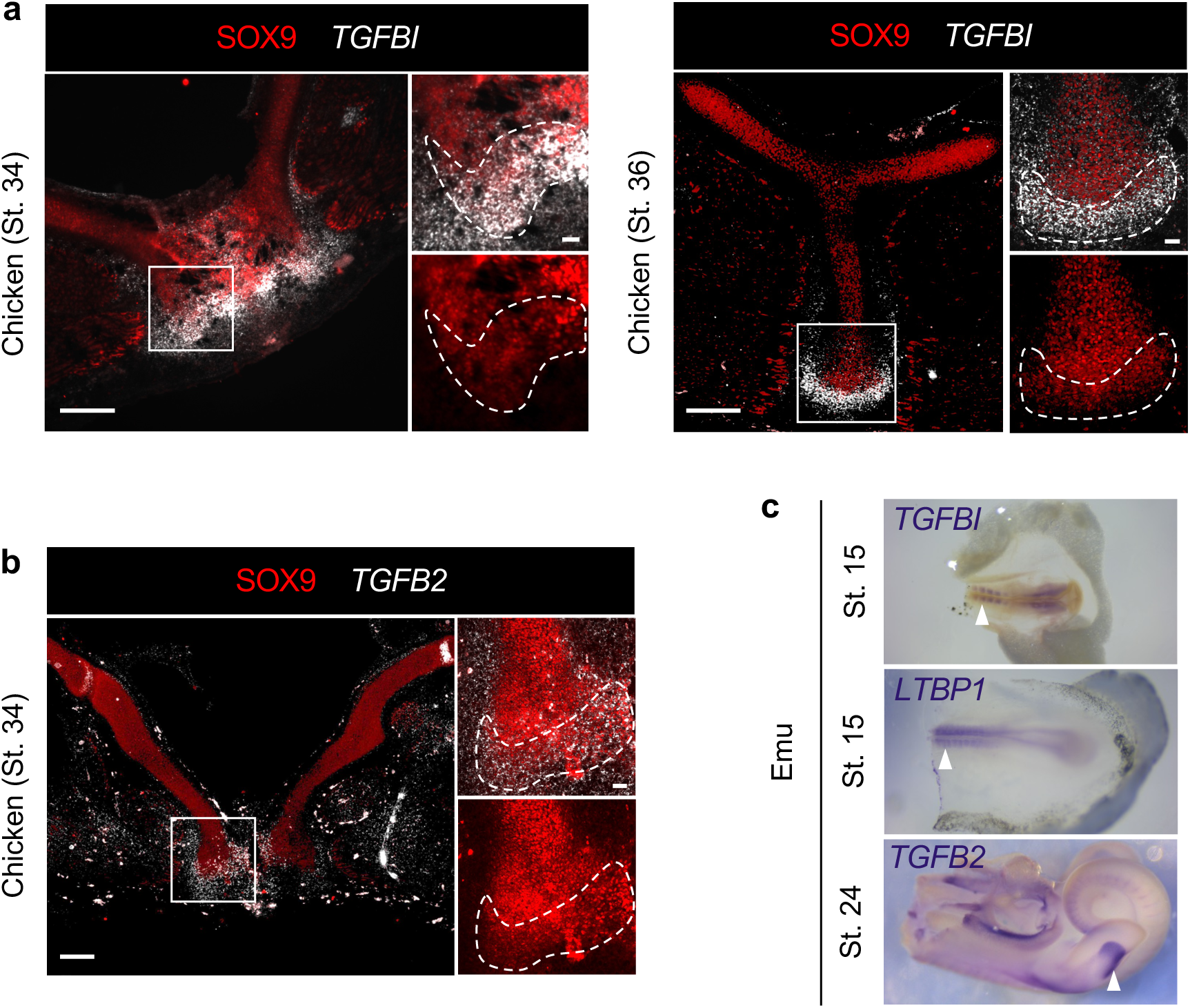
Colocalization of SOX9 with *TGFBI* and *TGF-β2* in chicken ventral SPs, and validation of emu riboprobes. **a** Detection of SOX9 (red) and *TGFBI* mRNA (white) in St. 34 and 36 chickens. In the dotted regions of the ventral parts of the sterna, the two signals are co-localized. **b** *TGF-β2* expression (white) is observed in SOX9-positive ventral SPs (dotted line). **c** Anti-sense riboprobes synthesized for emu *TGFBI*, *LTBP1*, and *TGF-β2* were validated by showing signals in somites at St. 15 (*TGFBI*, *LTBP1*) and in the limb mesenchyme at St. 24 (*TGF-β2*) (arrowheads), indicating that the probes worked normally. Scale bars, 200 μm; inset 30 μm.

**Supplementary Figure 6.**
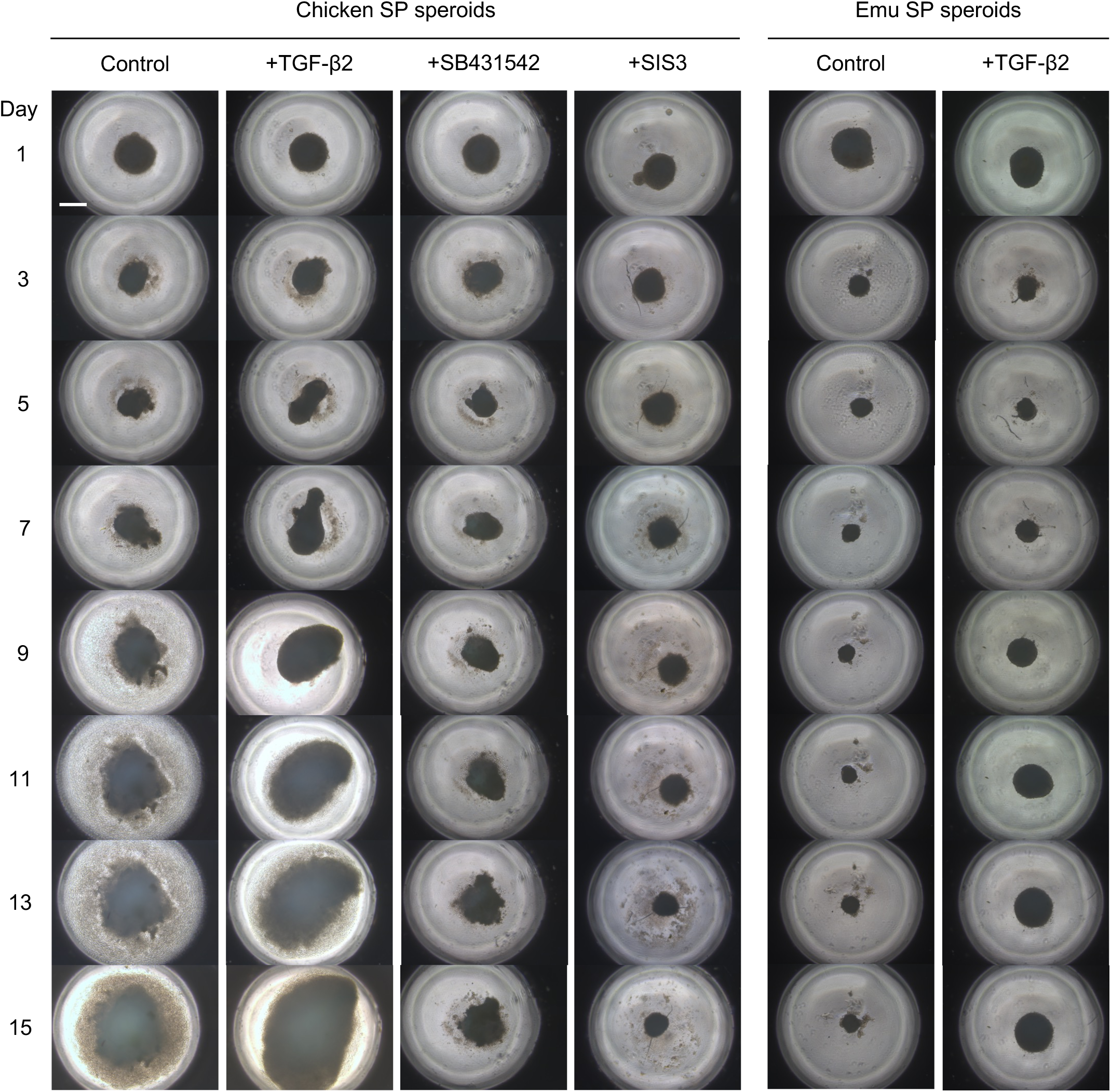
Chicken and emu SP spheroid cultures. Time course of spheroid morphological dynamics under each condition. Scale bar, 500 μm.

**Supplementary Figure 7.**
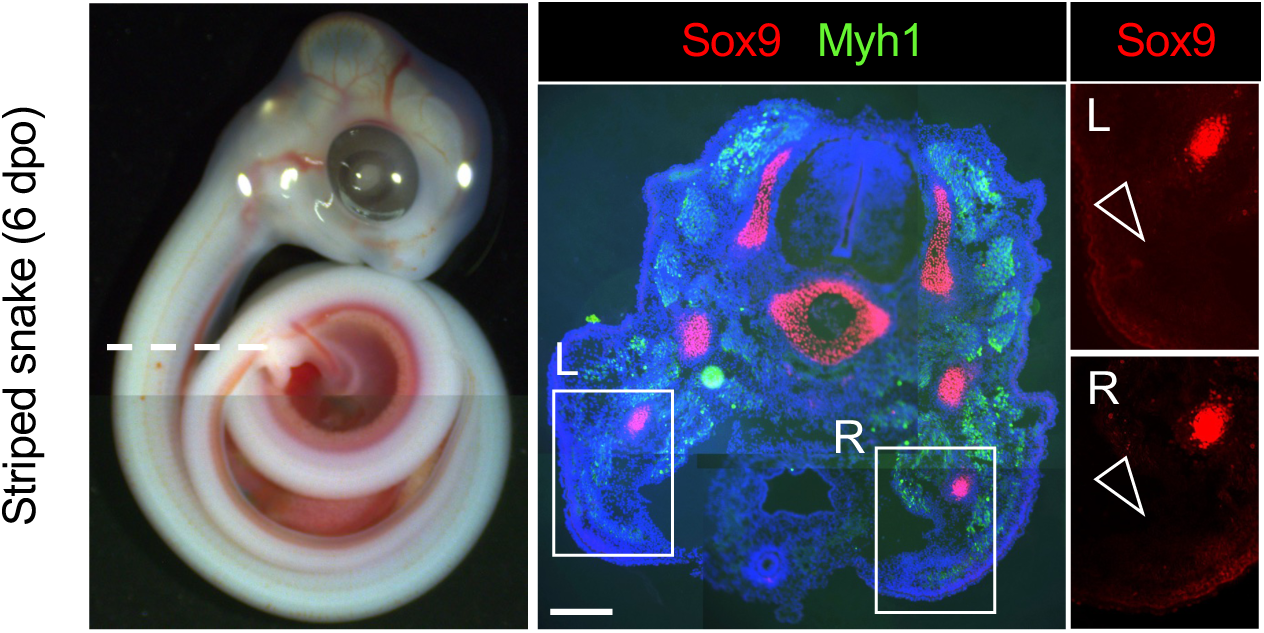
Absence of SOX9-positive SPs in the snake embryo. A transverse section of the anterior thoracic region (7-8^th^ somite level) of a Japanese striped snake embryo at 6 days post oviposition (6 DPO, developmentally comparable to chicken/emu St. 30), immunostained with antibodies against Sox9 (red) and Myh1 (cyan). Sox9 signals were absent in the ventral LPM regions on both sides (open arrowheads). Scale bar, 200 μm.

**Supplementary Movie 1**

**Three-dimensional rendering of an adult chicken sternum in orthographic view**

**Supplementary Movie 2**

**Three-dimensional rendering of an adult emu sternum in orthographic view**

## Notes

### Competing Interest Statement

The authors have declared no competing interest.

